# Microbe-metabolite associations linked to the rebounding murine gut microbiome post-colonization with vancomycin resistant *Enterococcus faecium*

**DOI:** 10.1101/849539

**Authors:** Andre Mu, Glen P. Carter, Lucy Li, Nicole S. Isles, Alison F. Vrbanac, James T. Morton, Alan K. Jarmusch, David P. De Souza, Vinod K. Narayana, Komal Kanojia, Brunda Nijagal, Malcolm J. McConville, Rob Knight, Benjamin P. Howden, Timothy P. Stinear

## Abstract

Vancomycin resistant *Enterococcus faecium* (VREfm) is an emerging antibiotic resistant pathogen. Strain-level investigations are beginning to reveal the molecular mechanisms used by VREfm to colonize regions of the human bowel. However, the role of commensal bacteria during VREfm colonization, in particular following antibiotic treatment, remains largely unknown. We employed amplicon 16S rRNA gene sequencing and metabolomics in a murine model system to try and investigate functional roles of the gut microbiome during VREfm colonization. First-order taxonomic shifts between Bacteroidetes and Tenerricutes within the gut microbial community composition were detected both in response to pretreatment using ceftriaxone, and to subsequent VREfm challenge. Using neural networking approaches to find co-occurrence profiles of bacteria and metabolites, we detected key metabolome features associated with butyric acid during and after VREfm colonization. These metabolite features were associated with *Bacteroides*, indicative of a transition towards a pre-antibiotic naïve microbiome. This study shows the impacts of antibiotics on the gut ecosystem, and the progression of the microbiome in response to colonisation with VREfm. Our results offer insights towards identifying potential non-antibiotic alternatives to eliminate VREfm through metabolic re-engineering to preferentially select for *Bacteroides*.

**Importance:** This study demonstrates the importance and power of linking bacterial composition profiling with metabolomics to find the interactions between commensal gut bacteria and a specific pathogen. Knowledge from this research will inform gut microbiome engineering strategies, with the aim of translating observations from animal models to human-relevant therapeutic applications.

## Introduction

Vancomycin-resistant *Enterococcus faecium* (VREfm) is a significant healthcare-associated pathogen. VREfm infections can be difficult to treat due to their intrinsic and acquired resistance to nearly all classes of antibiotics (1). The World Health Organisation categorizes VREfm as a “High Priority” bacterial pathogen, advocating research to stop the global increase in antibiotic resistance (2). Recent studies highlight the importance of the gut microbiota in modulating the growth and virulence of VRE*fm* in the gastrointestinal ecosystem. For instance, the depletion of normal gut flora using antibiotics exacerbates the severity of VREfm infection (3), whereas transplant of commensal species including a consortium of *Clostridium bolteae*, *Blautia producta*, *Blautia sartorii*, and *Parabacteroides distasonis* (4, 5) can drive established VREfm colonisation to below levels of culture detection. Specifically, *B. producta* – a colonizer of the colon – reduces VREfm growth *in vivo* by secreting a lantibiotic (6). These observations raise the intriguing possibility that metabolic traits act in concert between pathogen and select gut commensals to confer mutual benefits during pathogen persistence. These findings also highlight the greater risk posed to immunocompromised patients when colonized with VREfm. For instance, allogeneic hematopoietic cell transplantation patients have gastrointestinal tracts that are dominated by VREfm as a result of losing a large portion of the intestinal commensal microbiota upon receiving broad-spectrum antibiotics as pre-treatment(7). Hildebrand et al., (2018) discovered long-term ecological impacts to the gut microbiome, with strong bacterial species turnover, after ceftriaxone treatment in humans (8). Further, mice receiving broad spectrum antibiotics (combination of metronidazole, neomycin, and vancomycin) showed markedly increased VREfm colonization of the caecum and colon. The compromised intestinal innate immune defenses in these animals allowed proliferation of VREfm caused by the antibiotic exposure and subsequently reduced the expression of antimicrobial molecules produced by bacteria in the intestinal mucosa (9).

The problem with VREfm is further complicated by the fact that *Enterococci* are members of the gastrointestinal tract microbiota; a key reservoir of antimicrobial resistance (AMR) genes, and potentially facilitating gene transfer within the gut microbiome (10). For example, the *vanB* resistance gene was detected in human faecal specimens that did not contain culturable VRE; and instead, demonstrated that isolates carrying the resistance transposon are anaerobic commensal bacteria, *Eggerthella lenta* and *Clostridium innocuum*(11). Colonisation of, and persistence in, the gastrointestinal tract therefore presents as a key mechanism for *de novo* VRE and may lead to severe invasive disease.

This current study aimed to understand the impact of antibiotics on the murine gut microbiota, and the subsequent colonisation pattern of VREfm. To this extent, we designed a murine model time-series study that consisted of two main perturbative phases: (1) antibiotic pre-treatment with ceftriaxone, and (2) VREfm challenge. Our 16S rRNA gene profiling analyses highlighted a first-order shift in bacterial biodiversity composition across time, a second-order clustering of samples associated with the experimental phases, and the transition of the post-VREfm colonisation gut microbiota and its metabolome towards resembling an asymptomatic carriage-like microbiome phenotype. This research provides support for engineering the metabolic potential of the gut microbiome using, for example, prebiotics as a non-antibiotic alternative for treating multi-drug resistant bacterial infections.

## Results

### Experimental design

The following experimental design was developed to address the hypothesis that there are specific murine gut microbiome factors that facilitates VREfm colonization; three groups of three C57BL/6 mice (co-caged wildtype males) were monitored, and fecal samples collected, over a fourteen-day period with two intervention time-points; including (1) ceftriaxone treatment administered at 0.5g/L in drinking water across a two-day period, and (2) colonisation (via oral-gavage) with 1 × 10^6^ VREfm ST796 per mice post-antibiotic treatment at a single timepoint. Mice were housed in groups of five and samples collected from the same three mice to represent technical replicates per cage; herein, each group of co-housed mice will be referred to as Group A, Group B, and Group C. The remaining two mice per group were reserved for microbiological assays. Table 1 highlights samples and datasets collected.

**Table 1:**
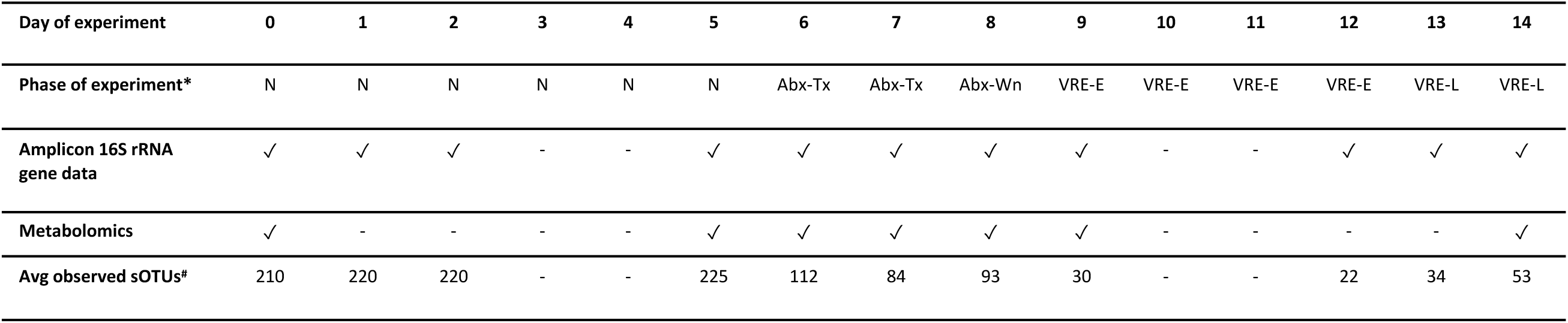
Summary of samples analyzed in this study Notes: *The key phases of the experiment where, N represents Naïve; Abx-Tx represents Antibiotic treatment; Abx-Wn represents Antibiotic weaning; VRE-E represents Early-phase post-VREfm colonisation; and VRE-L represents Late-phase post-VREfm colonization; #The average number of sOTUs observed across all mice for each day of the experiment; ‘✓’ Sample processed; ‘−’ Data unavailable

### Amplicon 16S rRNA gene sequencing revealed first order shifts in bacterial community composition

Amplicon 16S rRNA gene sequencing was performed to capture the bacterial community composition in an effort to track changes in response to antibiotic pre-treatment and VREfm colonization. Bacterial community profiles were assessed in fecal samples from nine mice before, during and after the two interventions (Table 1). A total of 71.32% of reads (10,519,073 reads) passed quality control, with 321,955 reads on average per sample and a total of 3574 exact variant sequence types (i.e., features), with an average of 118 features per sample, and an upper bound of 246 features (when rarefied to 20,000 reads). Alpha rarefaction analysis demonstrated sufficient sequencing depth to capture microbial diversity to saturation (Supplementary Figure 1).

The biodiversity profiles of each sample were compared and showed that key sOTUs were differentially abundant throughout the course of the experiment (Figure 1). There was a shift in the dominance of Bacteroidia (Bacteroidetes; light green coloured bars) during the naïve phase of the experiment to Mollicutes (Tenericutes; fuschia coloured bars) in response to ceftriaxone treatment, with a return to the predominance of Bacteroidia during the late-phase of the experiment, post-VREfm colonisation (i.e., days 12 – 14). Of note is the predominance of Lactobacillales in Mouse 1 – 3 from Group A (Figure 1).

**Figure 1.**
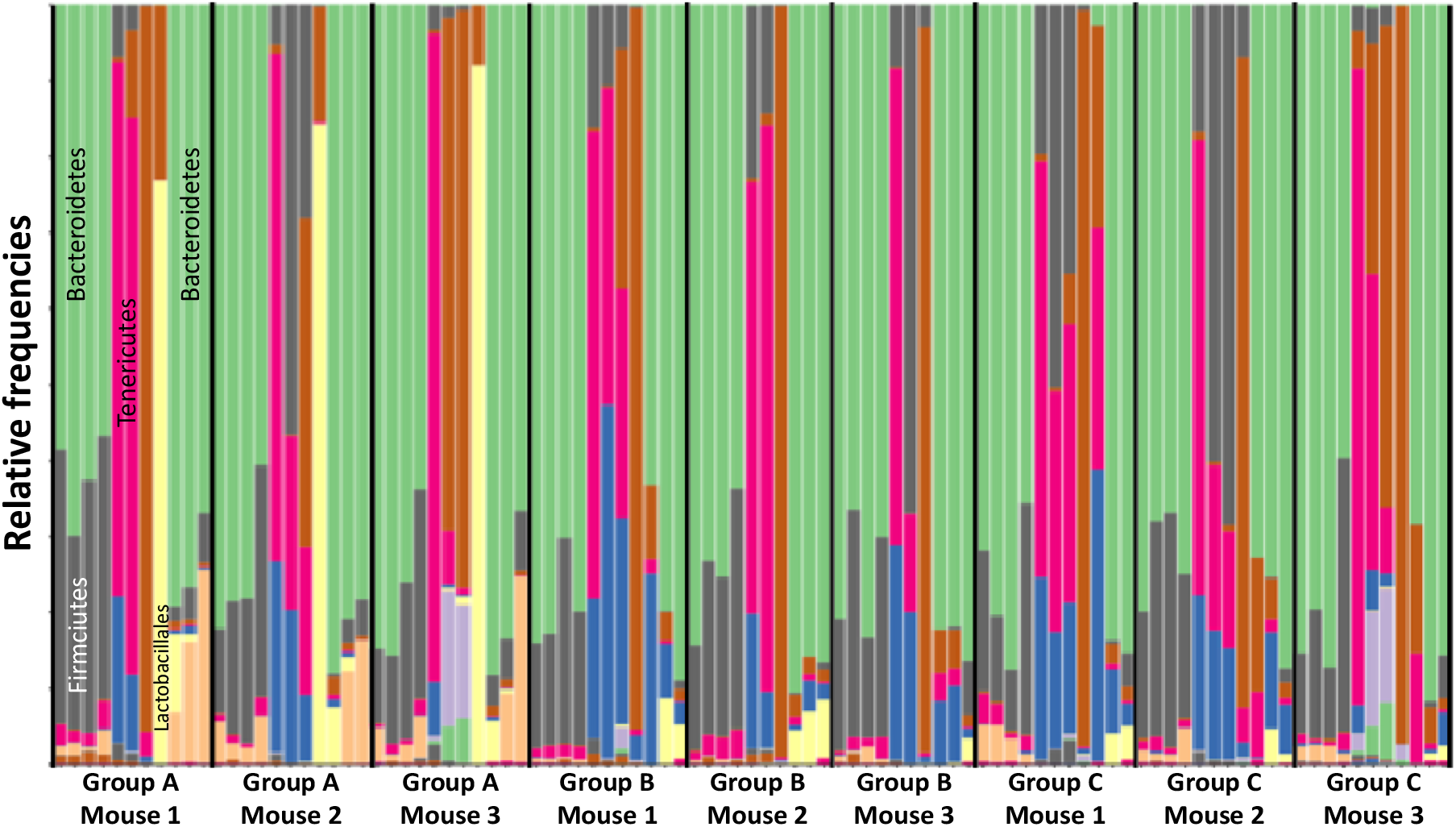
Biodiversity plot of sOTUs as relative frequencies at the taxonomic level of Class. First-order shifts in microbial community composition, as revealed by 16S rRNA gene community profiling, from a predominance of Bacteroidetes to Tenericutes and return to Bacteroidetes was observed. Each column displays the relative bacterial community composition in a mouse faecal sample collected daily and sorted by the chronology of the experiment (i.e., Day of experiment; Table 1). The columns are further sorted by group (i.e., Group A, Group B, and Group C) and individual mice within each group (mouse 1, mouse 2, and mouse 3). Stacked bars are presented as relative frequencies at the taxonomical level of Class, however, annotations of key taxa are at the phyla level (Bacteroidetes, green; Firmicutes, grey; and Tenericutes, fuschia) or Order level (Lactobacillales, yellow).

### The murine gut microbiota responds to antibiotics and microbial community richness begins to rebound three-days post-VRE colonisation

Principal coordinate analysis (PCoA) of the unweighted UniFrac (12) distances was used to assess clustering of fecal samples based on bacterial composition. This assessment showed that the fecal microbiota from samples collected from each phase clustered together but were clearly separated between phases; after exposure to ceftriaxone, and challenge with VREfm post-antibiotic treatment (Figure 2A). Permutation-based statistical testing demonstrates the groups are significantly different from one another (Supplementary Figure 2). Temporal tracking of the changing microbiomes against each mouse on the PCoA sample-space demonstrated a clear, unidirectional trajectory that followed the chronology of the experiment (animation as a supplementary file). Procrustes analyses of weighted and unweighted UniFrac distances showed that the same general patterns on the sample space were preserved, meaning that there is congruency in global spatial patterns between qualitative and quantitative measures of community dissimilarity (Supplementary Figure 3).

**Figure 2.**
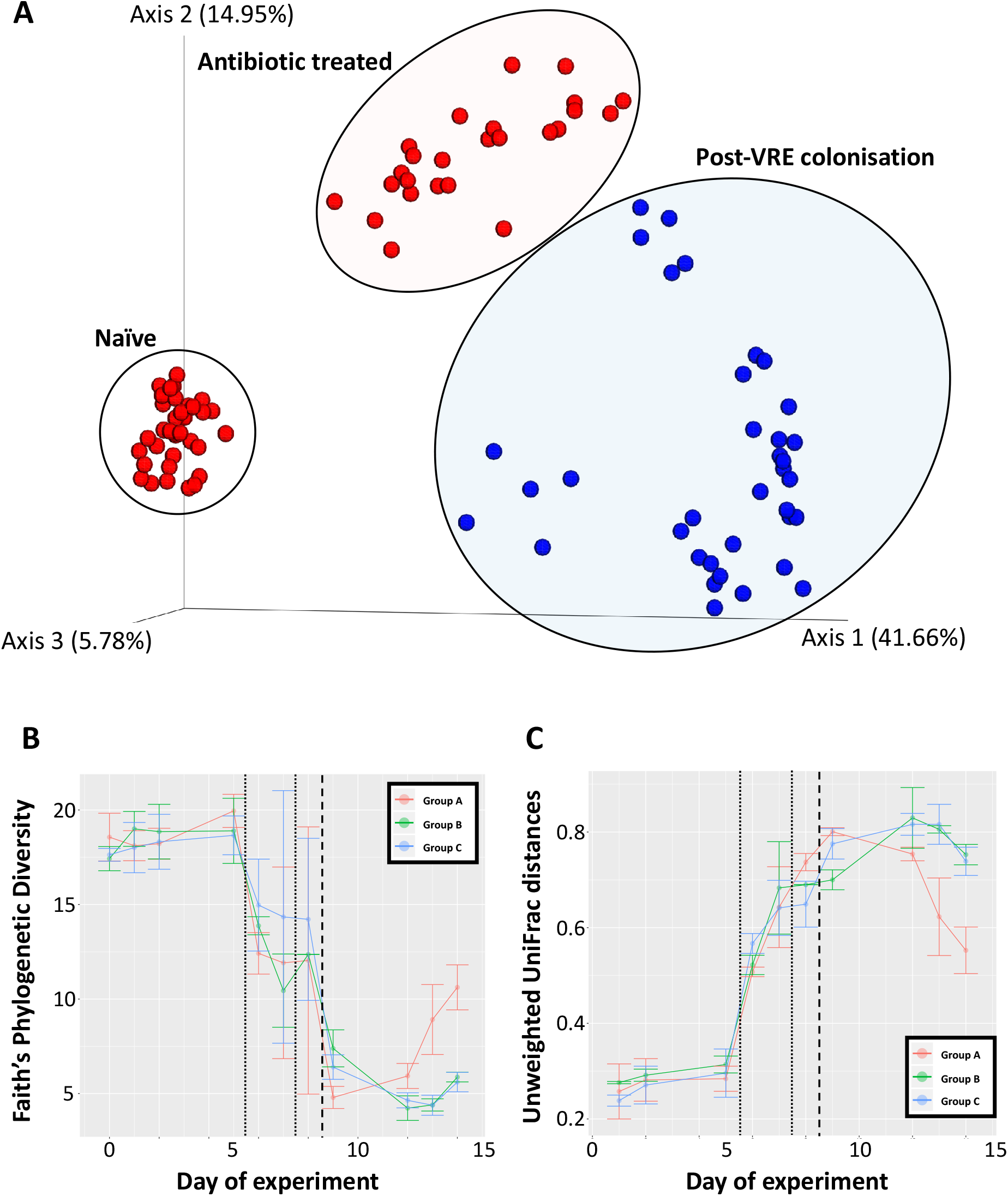
Diversity analyses. **(A)** Principal coordinate analysis plot of unweighted UniFrac distances. Data points are projected onto the sample space, and coloured by pre-VREfm colonisation (red), and post-VREfm colonisation (blue). *n.b.,* circles and ellipses function to highlight the separation of experimental phases and do not indicate statistical confidence intervals. Principal coordinate axis 1 explains 41.66% of the variation observed between the naïve microbiota and those from the post-VREfm colonisation phase. **(B)** Community richness of the murine gut microbiome, as measured by Faith’s Phylogenetic Diversity, in response to ceftriaxone treatment and challenge with VREfm; **(C)** community dissimilarity distances, as calculated by unweighted UniFrac, of each time point relative to day 0 (naïve phase).

Analysis of community diversity (Faith’s Phylogenetic Diversity Index) revealed a stable and rich microbial community during the naïve phase preceding a sharp decrease following antibiotic treatment, and a further decrease immediately following VREfm colonisation (Figure 2B). Of note is the responsiveness of the microbiota (within 24 h) to the removal of antibiotics at the end of day 7. Community richness began to rebound at approximately three-days post-VREfm colonisation (i.e., day 12), with Group A demonstrating a faster rate of rebound compared to Group B and C. Calculating the distances of dissimilarity (unweighted UniFrac distances) of each mouse microbiota timepoint relative to Day 0 (a proxy for the naive bacterial community phenotype) revealed a small dissimilarity distance for samples collected during the naïve phase, and an increasing dissimilarity distance following antibiotic treatment (day 6) and VREfm colonisation (day 9; Figure 2C). There was a downward trajectory in distance scores at three-days post-VREfm colonisation (i.e., day 12); Group A followed a sharper return to a microbiota resembling day 0. These observations suggest that mice were transitioning towards a persistent carrier-like state, and that the rebounding community richness towards levels representative of the naïve phase was by a microbial community structure that resembled the naïve phase. Additional studies where the time-frame of post-VREfm challenge extends beyond one-week of monitoring are needed to understand if the perturbed microbiome will return to resemble an absolute naïve state or arrive at a new, altered state.

### Multinomial regression identifies sOTUs most positively associated with VREfm colonisation

Multinomial regression using *Songbird* was employed to identify sOTUs that were most positively, and negatively, associated with the post-VREfm colonisation phase (Figure 3). The five most positively associated sOTUs were *Enterococcus*, *Bacteroides*, *Erysipelotrichaceae*, *Catabacter*, and *Lachnospiraceae*; while the five most negatively associated were *Clostridiales*, *Adlercreutzia*, *Mollicutes*, *Peptostreptococcaceae*, and *Clostridiales*. Temporal tracking of exact sequence variants (ESV) demonstrated that the ESV feature classified as *Enterococcus* - and identified as the most positively associated with the post-VREfm colonisation phase - was most abundant on days of challenge; confirming that this ESV likely was the ST796 VREfm colonization challenge organisms (Figure 3). There were a further eight ESV features classified as *Enterococcus*, however, they were absent during the days representing VREfm colonisation, and lacked positive associations with the post-VREfm colonisation phase, suggesting that these features represent murine gut commensal Enterococci.

**Figure 3.**
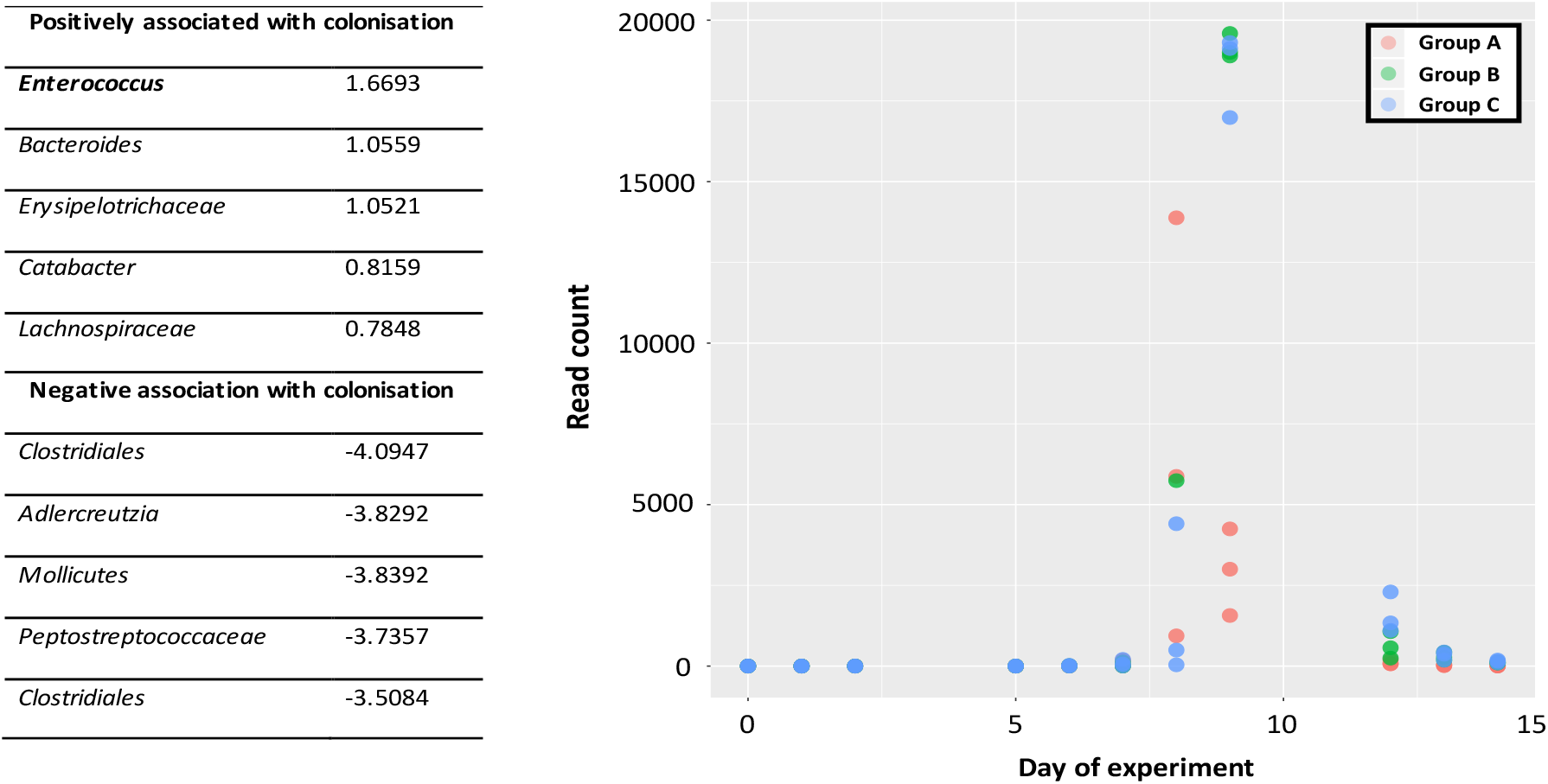
Multinomial regression. Multinomial regression identified an *Enterococci* exact sequence variant as the most positively associated with the colonisation phase (log-fold change score of 1.6693). Read counts for the *Enterococci*-ESV tracked daily across the experiment showing high abundance during days of VREfm challenge.

### Molecular networking identifies differential metabolome profiles

Duplicate fecal samples from key timepoints throughout the experiment (i.e., days 0, 5, 6, 7, 8, 9, and 14) were analysed by data dependent tandem mass spectrometry (MS/MS) performed on a liquid-chromatography quadrupole time-of-flight (LC-QTOF) to monitor changes in the murine gut metabolome (Table 1). Polar metabolite analysis was preferenced in an effort to broadly capture primary metabolites which play a key role in “metabolic hand-offs” that define interspecies interactions. Analysis of the global metabolome profile of each sample was measured based on their overlapping molecules, and a PCoA plot using a binary Jaccard distance metric through the Global Natural Products Social Molecular Networking (GNPS) platform (13). A separation of metabolite profiles along PCoA1 was observed (57.34% Figure 4A). Metabolomes from the naïve, and late-VREfm colonisation phase tended to cluster together, while samples from the post-antibiotic phases including the early-VREfm colonisation phase clustered together. Supporting pairwise PERMANOVA testing (Supplementary Figure 4) highlights naïve and early-VRE samples are significantly different, while late-VRE has a lower distance to naïve compared to antibiotic treated, antibiotic wean and early-VRE samples.

**Figure 4.**
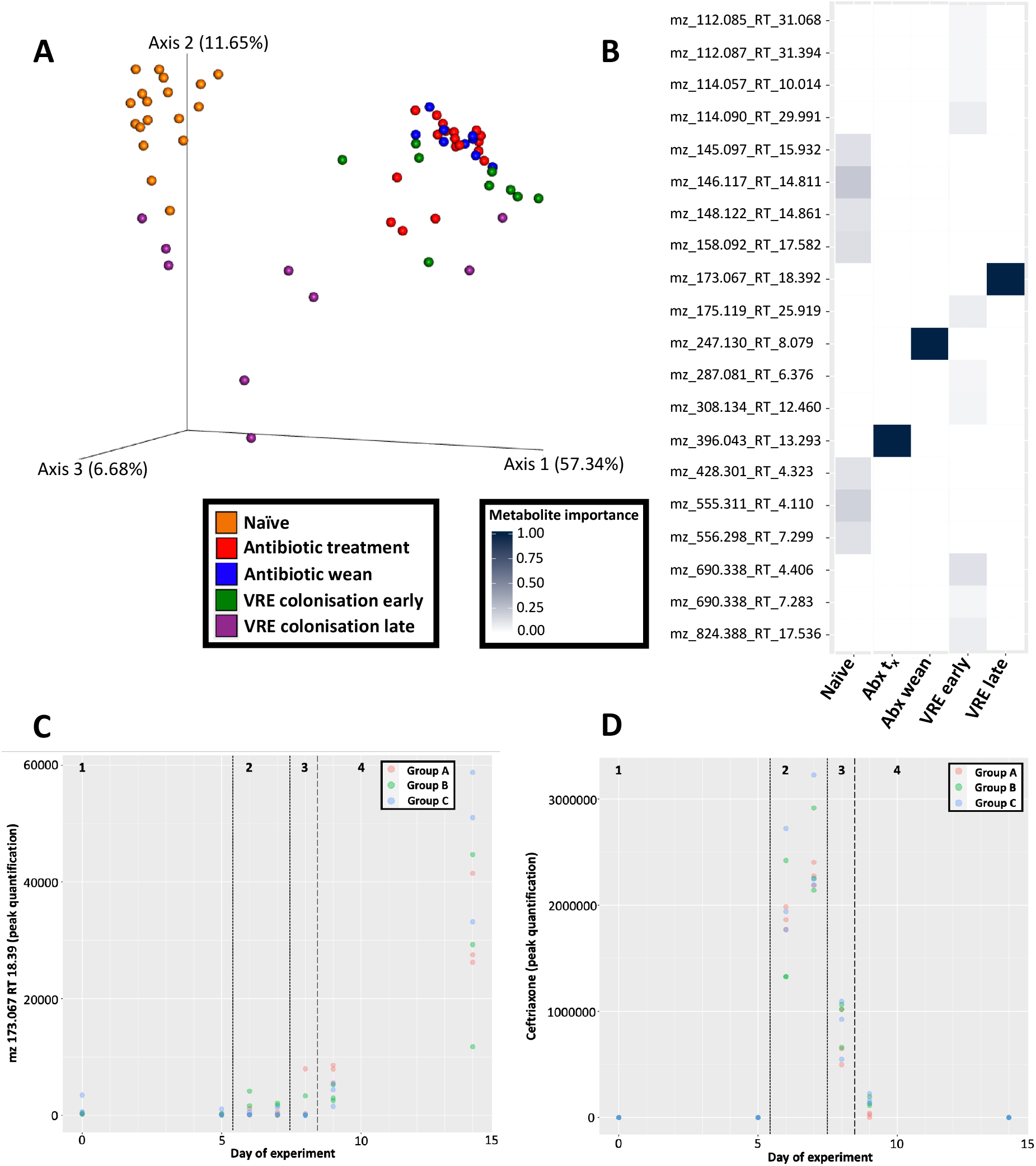
Metabolomic analyses. **(A)** Emperor plot displaying principal coordinate analysis of binary Jaccard distances of metabolomic profiles. Samples represent (colour coded) the naïve (orange), antibiotic-treatment (red), antibiotic-weaning (blue), early-VRE colonisation (green), and late-VREfm colonisation (purple) phases. **(B)** Random Forest classifier identifying metabolite features (spectra) for each phase of the experiment. Heatmap is colour-coded from low ranking score (white; i.e., lowest importance) to high ranking score (dark blue; highest importance). Metabolite features are labelled by their mass-charge ratios and retention times for the reason that current databases do not capture their chemical structure and/or identifications. **(C)** Peak quantification values for feature 6325 (m/z= 172.0671 RT = 18.39) present in abundance during *VRE colonisation late* (phase 4). **(D)** Peak quantification values for ceftriaxone (m/z = 555.0537 RT = 13.30) tracked across the experiment. Ceftriaxone values are highest during *antibiotic treatment* (phase 2) and begins to wane during *antibiotic wean* (phase 3).

Random forest analysis of spectra profiles from LC-MS/MS was used to predict experimental phase and rank the importance of metabolite association with each experimental phase. The top metabolite features for each experimental phase are highlighted in Figure 4B. Unique profiles of metabolite features were observed for each phase of the experiment. Importantly, the late-VREfm colonisation phase captures an unknown metabolite (feature 6325) with a mass-to-charge ratio (m/z) of 173.067 and retention time (RT) of 18.392; this metabolite is exclusively present during what represents the *transition* towards resembling the naïve microbiome. Manual curation of feature 6325 in positive ion mode predicts a molecular formula C_5_H_8_N_4_O_3_ with ≥ 10 ppm in mass error. The major peaks in the MS/MS spectrum for feature 6325 are: 173.07 (precursor ion, [M+H]+ assumed), 155.06 (precursor ion, 18(H_2_O), 113.05 (155.06 product ion, 42.01 (likely C_2_H_2_O), 43.03 ([C_2_H_2_O+H] + product ion further supporting neutral loss of C_2_H_2_O); given the summation of results, the chemical structure of feature 6325 is likely to contain a N-acetylated hydroxyl group. Peak quantification values indicate its presence during the late phase of VRE colonisation (Figure 4C). Further manual curation of MS/MS data identified ceftriaxone as feature 3901 with an m/z = 555.0537 and RT = 13.30, and mostly abundant during days of antibiotic exposure (Figure 4D).

### Bacteroidales-associated metabolites implicated in late-phase post-VREfm colonisation

A distinct profile shift in microbe and metabolite abundances (as calculated by multinomial regression) was observed, particularly during late-phase VREfm colonisation (Supplementary Figure 6). Shallow neural networking analysis with *mmvec* was used to predict microbe-metabolite interactions through their co-occurrence probabilities (Figure 5). Sequential biplots captured the shift in experimental phases and highlighted the co-occurrences of microbiota and metabolomic datasets (Fig. 5A,B,C). There was a strong *Enterococcus* effect as indicated by the magnitude of the corresponding arrow, and the rebounding species during the late-phase VREfm colonisation are predominantly *Bacteroidales* sOTUs (Figure 5C) with co-occurring metabolite features m/z 173.067 RT 18.392, and m/z 167.083 RT 25.277. Metabolite feature mz 245.055 RT 7.945 was ranked as being highly associated with the post-VRE colonisation phase. These results integrate microbial and metabolite datasets to reveal which microbes may be responsible for detected metabolites. In this instance, the metabolite present during the phase representing a *transition* towards a microbiome approximating the naïve state (feature 6325 m/z 173.067 and RT 18.392; Figure 4B) is linked with *Bacteroidales* (Figure 5A).

**Figure 5.**
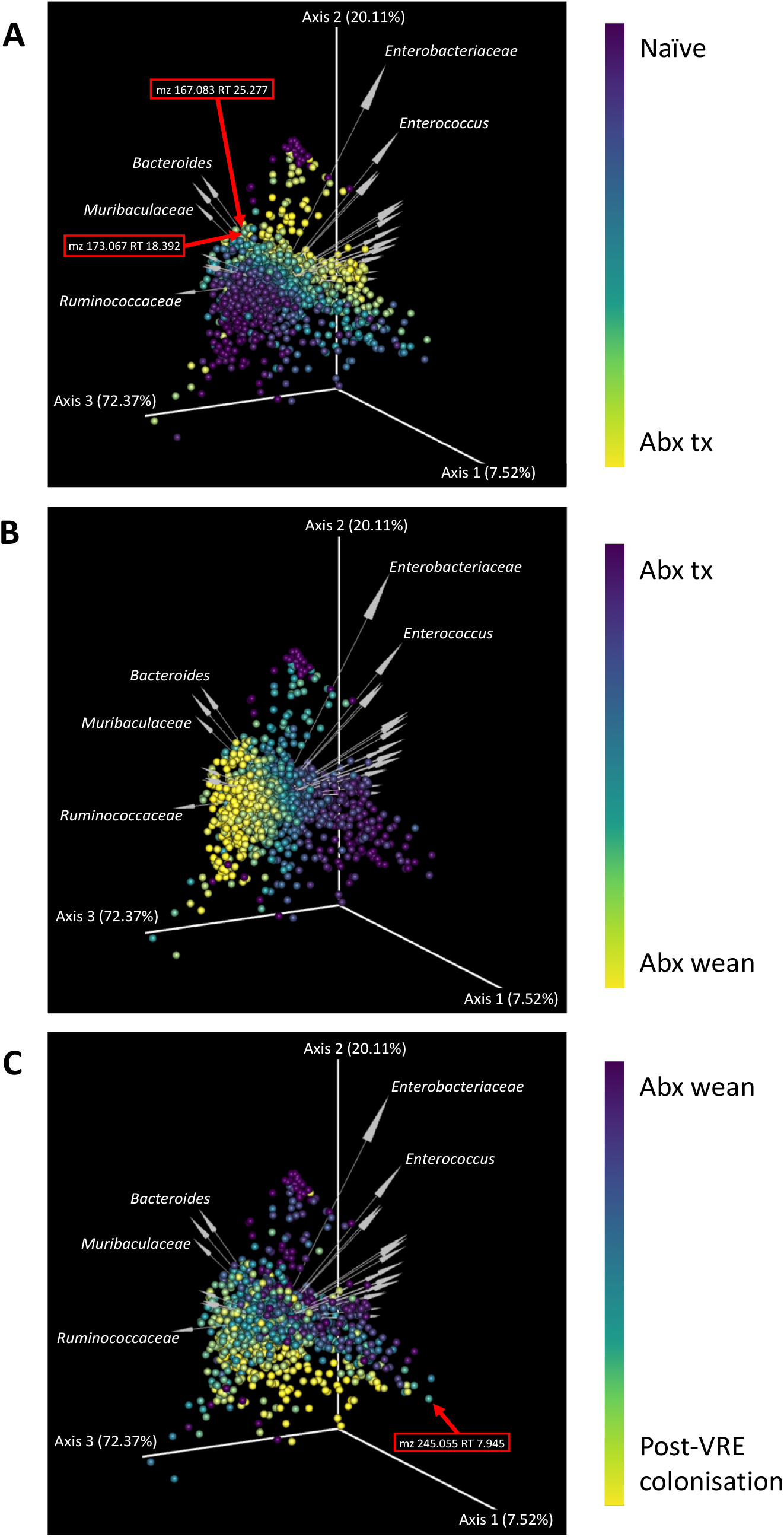
Microbe-metabolite vector biplots. Sequential biplots highlighting the changing metabolite differentials across each key phase of the experiment; where *abx tx* corresponds to the *antibiotic treatment* phase, and *abx wean* corresponds to the period where antibiotics were removed for a 24-hour period prior to colonisation with VREfm. Each point on the sample space represents metabolites, and arrows represent microbes. Microbe and metabolite features are fixed upon the sample space, with gradient coloring of metabolites indicating the transition across key phases of the experiment. The distance between each point is indicative of metabolite co-occurrence frequency, and the angle between arrows indicate microbial co-occurrence. The directionality of the arrows describes the variation in the metabolites explained the microbes represented by the arrows. For example, metabolite feature 6325 (m/z 173.067 RT 18.392) is demonstrated to co-occur with *Bacteroides*. Information about the abundances of these co-occurring features are provided as heatmaps in Supplementary Figure 6.

## Discussion

In this study of the murine gut ecosystem, we employed a mouse model of gastrointestinal tract colonisation that replicates the shift in bacterial composition when patients enter the healthcare system, develop an imbalance in microbiome as a result of pre-treatment (e.g., antibiotic treatment), and are subsequently colonised with a hospital superbug (14). The resolution of current studies describes a consortium of commensal microbes that can, for example, reduce the magnitude of VREfm colonisation (4, 6); however, understanding the key metabolic shifts relative to the gut microbiota remains challenging (15). Here, we employed amplicon 16S rRNA gene sequencing and high-resolution mass spectrometry metabolomics in an effort towards determining microbiota-metabolome interactions during VREfm colonisation. We demonstrated clear changes in the gut microbiome in response to ceftriaxone and VREfm challenge.

Conceptual and statistical advances in analysis of amplicon 16S rRNA gene data (16, 17) whereby OTUs are clustered at a 99% nucleotide similarity threshold allows for the identification of exact sequence variants (ESV). Query against an error-corrected database (17) can detect multiple ESVs that may be classified to the same taxonomic rank. For example, our analyses identified multiple ESVs classified as *Enterococci*; however, when the relative abundances were tracked across the chronology of the experiment, only one *Enterococci*-ESV was dominant in relative abundances and most positively associated with the days of post-VREfm challenge (Figure 3). This highlights the resolving power to differentiate between commensal and pathogenic strains of *Enterococci* when the composition of the microbial community is considered. The fact that this was achievable at the level of amplicon 16S rRNA gene sequencing alludes to the possibility of implementing microbiota screenings as routine diagnostics for patients entering healthcare systems. Further, first-order level shifts in microbial community composition was observed in response to ceftriaxone and subsequent VREfm challenge (Figure 1). At three days post-VREfm colonisation (i.e., day 12), the microbiome richness begins to rebound; suggesting that mice are transitioning towards a persistent carrier-like state. Interestingly, Group A cohort exhibited a faster rate of rebound that may be facilitated by their initially higher microbial community richness and predominance of Lactobacillales during day of VREfm challenge (Figure 2B); this observation supports the need to prescreen “baseline” microbiota profiles of patients upon admission into hospital for the reason that it is not necessarily which microbial populations are removed post-perturbation (e.g., antibiotic pre-treatment) but instead, which populations persist that drives the responding phenotype. We can begin to assess patients from across different wards (e.g., intensive care unit, oncology, neurology, and healthy cohorts), and build a database of microbiome profiles that can be used as biomarkers to predict: (1) the susceptibility of patients to develop persistent bacterial colonisation, and (2) propensity to clear the pathogen once colonized. The clinical implication is that new patients are screened and identified (via beta-diversity meta-analyses) by these biomarkers and placed in bedding cohorts accordingly; thereby improving infectious disease management and isolation precautions within health-care associated ecosystems.

The shortlist of microbes ranked as most negatively associated with the colonisation phase (*Clostridiales*, *Adlercreutzia*, *Mollicutes*, *Peptostreptococcaceae*, and *Clostridiales*; Figure 3) are hypothesized to play a role in maintaining the health of the animals. Indeed, amongst the microbes identified, are known short chain fatty acid (e.g., butyric acid) producers (18, 19) that supports, and expands upon those previously identified by Caballero et al., (2017). Further, the use of Deblur to identify ESVs facilitates the temporal tracking of their relative abundances to inform selection of primary fecal samples that will provide the best probability (i.e., highest relative abundance) of culturing target taxa for downstream screening of probiotic potential. However, translating animal-derived observations from experimental animal models to human clinical situations remains challenging; particularly where the key microbes are rodent-specific. One solution may be to integrate metabolomics to reveal shared metabolic capacity amongst taxonomically divergent microbes. Our supervised classifying approaches suggests an altered metabolome composition during the late-phase of VREfm challenge that may facilitate the apparent “suppression” of VREfm to levels below the limit-of-detection by culture. Despite the caveat of poor resolution in current databases to link metabolite features to associated chemical structures, microbe-metabolite vector analysis linked metabolite feature 6325 (m/z = 173.067 RT = 18.392) to *Bacteroides* (Figure 5). Our efforts towards manually identifying feature 6325 suggests a chemical formula of C_5_H_8_N_4_O_3_ and a structure likely to contain a N-acetylated hydroxyl group; a putative annotation (through *pubchem* search) is 3-hydroxy-4-(nitrosocyanamido) butyramide. Butyramide is the amide group of butyric acid, a short-chain fatty acid that has been shown to play a key role in colonisation resistance against intestinal pathogens (20–23). Further research to comprehensively characterize interactions between microbe and metabolites will be critical to address the gaps in our understanding of the biochemical parameters that define interspecies microbiome interactions during antibiotic pre-treatment and persistent infections.

The resolution of our results provides the basis in which to begin to identify non-antibiotic alternatives to engineer the gut microbiome through prebiotic interventions (e.g., butyric acid), and translating animal studies to human-relevant therapeutic applications by delineating taxonomically diverse microbes with shared metabolic capacity. Here, achieving integrative omics to link microbe-metabolite associations, our findings adds support to the incorporation a microbiome profiling approaches into routine clinical microbiology, particularly in the context of monitoring the impacts of antibiotic use.

## Methods

### Mouse gastrointestinal colonization model

Six-week old wild type C57BL/6 male mice were used to establish an animal model of gastrointestinal colonisation with VREfm. Mice were co-housed, they have free access to food (ordinary chow) and water and have environmental enrichment (e.g., fun tunnels, chew blocks, and tissue paper). The light/dark cycle was 12 h light/ 12 h dark, and cages were changed weekly. Mice were pretreated with 0.5 g/L ceftriaxone in drinking water for two-days followed by an antibiotic wean period of 24 h. Mice were then challenged with 1 × 10^6^ cfu VREfm ST796.

### Genomic DNA extraction and sequencing

Whole community gDNA was extracted from mouse faecal samples using the Qiagen PowerSoil DNA Extraction Kit (formerly MoBio) following the manufacturer’s protocol. A preprocessing step of mechanical lysis was incorporated using a Bertin Technologies Precellysis 24 machine for one round of a 40 second-cycle at 6000 rpm. The V4 region of the bacterial 16S rRNA gene was amplified using small subunit 515 forward Golay-barcoded, and SSU806 reverse primers following the Earth Microbiome Project protocol (www.earthmicrobiome.org;(24)), and sequenced using the Illumina MiSeq platform (V2, 300 cycles; Illumina Inc., San Diego, USA). Sample metadata and sequence data for the current study were deposited into *QIITA* (25) - an open-source management platform for research studies with multiple ‘omic datasets - under the project title *Mouse-hospital VRE colonization with ceftriaxone treatment* (Study ID 11737). Further, primary derived data (e.g., BIOM tables) used to produce results can be found within QIITA study ID 11737. Raw sequence data are also available through the NCBI SRA, BioProject number: PRJNA590613

### Amplicon 16S rRNA gene profiling analyses

Sequence data were processed within the *QIITA* (v0.1.0) framework for quality control (split libraries v. q2.1.9.1), demultiplexing, trimming sequence reads to a length of 150 nt, and picking sub-operational taxonomic units (sOTUs) using *Deblur* (v1.1.0) to resolve single-nucleotide community sequencing patterns (*i.e*. feature identification of sOTUs;(17)). The output BIOM files were further processed using *QIIME2* (v2019.7) for downstream statistical analyses(26). Alpha rarefaction curves were generated to determine if each sample had been sequenced to saturation; the feature table was subsequently rarefied to 20,000 reads per sample. Taxonomy was assigned using the *sklearn* classifier (27) and *Greengenes 13.8 99% OTUs from 515F/806R region of sequences* classifier available from https://docs.qiime2.org/2018.4/data-resources/. Furthermore, relative abundances of each taxa were visualized as bar plots using the QIIME2 *taxa* plugin. A phylogenetic tree was constructed using fragment-insertion (*QIIME fragment-insertion sepp;* (28)) to guide phylogenetic-aware statistical analyses generated using the *QIIME2* plugin, *q2-diversity core-metrics-phylogenetic*; key metrics computed by default include both alpha-diversity (e.g., Shannon’s diversity index, Faith’s phylogenetic diversity, and Evenness), and beta-diversity (e.g., Bray-Curtis distance, and unweighted UniFrac distances) metrics. The unweighted UniFrac distance matrix (12) was used to compute first distances and calculate distances relative to Day 0 as baseline between sequential states (*QIIIME longitudinal first-distances*); *ggplot2* (R v3.6.0, https://ggplot2.tidyverse.org) was used to visualize the distance scores as line plots. *Emperor* was used to visualize principal coordinate analysis plots of unweighted UniFrac distances. Permutation-based statistical testing (PERMANOVA) on unweighted UniFrac distances was used to determine if samples grouped by *phase of experiment* were significantly different from one another (*q2-beta-group-signifcance*). *Songbird* (https://github.com/mortonjt/songbird) was employed to determine the importance (*i.e.,* fold change) of each sOTU in relation to a given metadata variable (e.g., VREfm-colonisation). Microbial features from all samples were split into training and test sets for supervised learning classifier analyses; 20% of input samples were allocated to train the Random Forest Classifier within QIIME2, the estimator method used for sample prediction. The different experimental phases were the response variables, while the 16S rRNA gene data were the features.

### Metabolite extraction and liquid chromatography tandem mass-spectrometry analysis

Duplicate fecal samples, as outlined in Table 1, were processed for polar metabolite extraction and analysis (days 0, 5, 6, 7, 8, 9, and 14). Feces were metabolically arrested by immediate collection into dry ice, and stored at −80°C until further processing. Metabolite extraction from the fecal samples was undertaken by the addition of 500 μL per sample of methanol:water solution (3:1 v/v) containing 2μM ^13^C-sorbitol and 8μM ^13^C,^15^N-valine, and 2μM ^13^C-leucine as internal standards. Fecal samples were homogenized at 1200 rpm for 30 minutes in a thermomixer maintained at 4°C, and mechanically disrupted and incubated for a further 15 minutes in the thermomixer. Samples were randomized for metabolite extraction.

Metabolite analysis of the extracted samples, pooled biological quality control (PBQC) samples, and 13 mixtures of authentic standards mixes was performed by liquid chromatography mass spectrometry (LC-MS) using hydrophilic interaction column (ZIC-pHILIC) and high-resolution Agilent 6545 series quadrupole time-of-flight mass spectrometry (QTOF MS) as described previously (29). PCoA of binary-Jaccard distances of test, standard mixes, and PBQC samples are presented in Supplementary Figure 5. Ions were analyzed in positive mode with full scan range of 85 to 1200 m/z and in data-dependent tandem MS mode to facilitate downstream metabolite identification. Mass spectrometry data were deposited into *QIITA* (25) under the project title *Mouse-hospital VRE colonization with ceftriaxone treatment* (Study ID 11737).

### Metabolomic analyses

The data-dependent tandem MS data was processed using *MZmine2* (v2.39) (30) to generate tabular matrices of metabolite features (i.e., m/z and retention time). Masses were detected, and chromatograms built using *Peak Detection* methods within *MZmine2.* Chromatograms were deconvoluted, and isotopic peaks grouped; grouped peaks were aligned using *join aligner*. *Peak list rows filter* method was applied to the aligned peaks, and gaps filled using *peak finder*. The following MZmine2 settings were applied for spectral processing; MS1 mass detection: 1E3; MS2 mass detection: 1E2; time span: 0.02; minimum height: 3E3; m/z tolerance: 10 ppm; pairing m/z range for MS2: 0.1; RT range for MS2 scan: 2 min; minimum peak height: 7E3; peak duration: 0.02 – 5.00, baseline: 5E3;0.001, or maximum chance: 2; Join Aligner: 75% 25% ratio split; intensity tolerance: 10%; and m/z tolerance: 5 ppm. Feature finding produced a data matrix of MS1 features and associated peak areas. Feature-based molecular networking outputs (*quant.csv*) were generated from *MZmine2* using the *export to GNPS* module, which contains MS1 feature information and a corresponding *mgf* file containing MS2 information linked to MS1 features. Metabolomic features were further analysed within the *Global Natural Products Social Networking (GNPS* v1.2.5; (13)) framework (UCSD, CA, USA). Tandem MS data were processed for identification, dereplication and quantification, including spectral library searches. For example, MS2 spectra of the unknown metabolites are compared with a library of MS/MS spectra generated from structurally characterized metabolites. Further information on the GNPS workflow, and molecular networking can be found in Ming *et al.,* 2016 (13). Further, manual interpretation – including, for example, determining molecular formula of the chemical in neutral charge structure, determining theoretical monoisotopic mass, and determining likely adduct - of MS/MS data was applied to identify unknown features.

### Neural networking to predict microbe-metabolite interactions

Microbiota and metabolome feature tables were analyzed using *MMVEC* ((31)https://github.com/biocore/mmvec) to identify microbe-metabolite interactions through their co-occurrence probabilities as predicted by neural networking. Conditional biplots were generated using *Emperor*. Further, microbe abundances and metabolite log centered abundance heatmaps were generated using primary derived data from multinomial regression analyses using *Songbird* (https://github.com/mortonjt/songbird).

## Acknowledgements

We thank the team at Metabolomics Australia (Bio21) mass spectrometry services. We thank Dr. Daniel Petras (UCSD) for helpful discussions.

## Supplementary figures

**Supplementary Figure 1.**
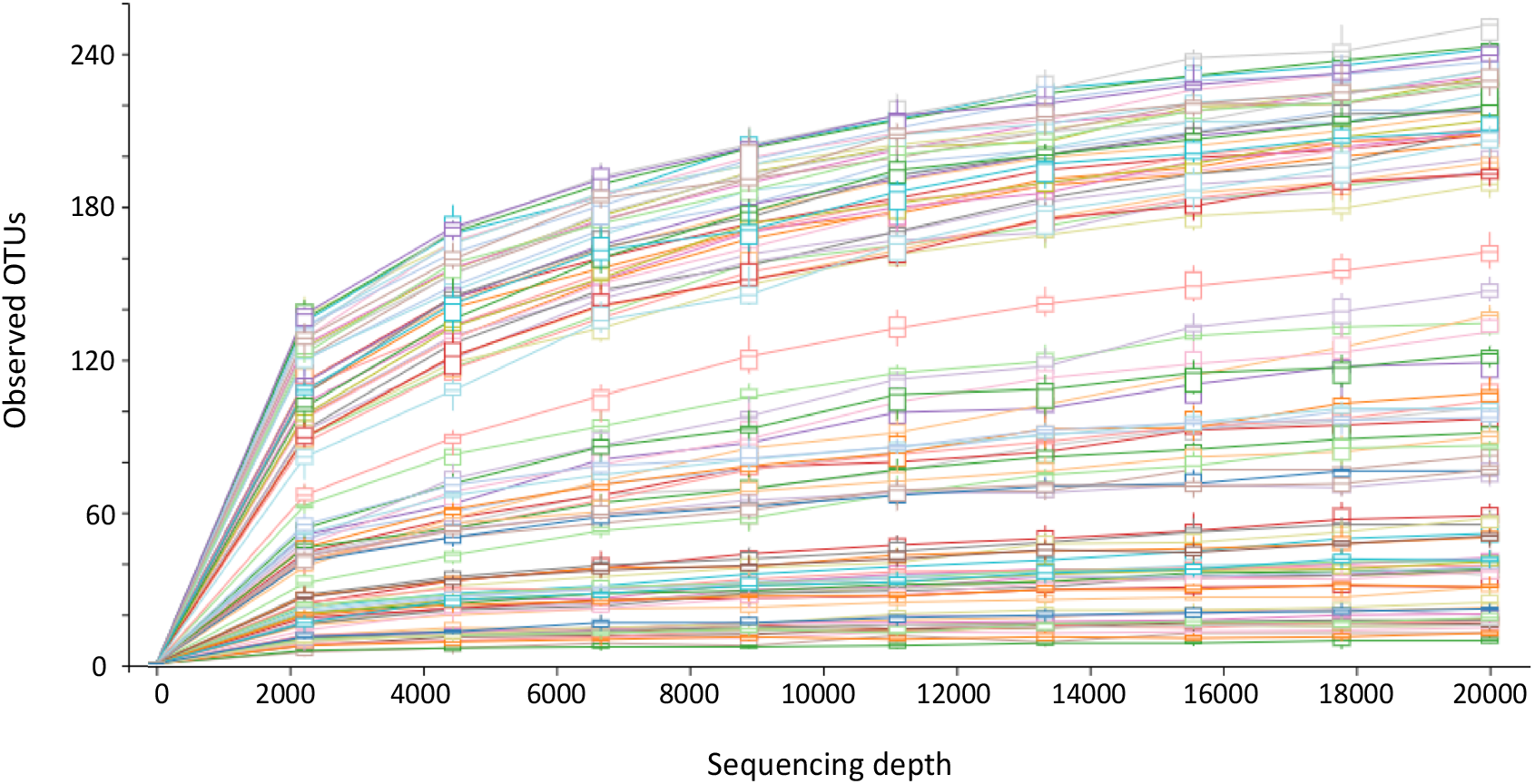
Alpha rarefaction plot of all samples rarefied to 20,000 reads per sample.

**Supplementary Figure 2.**
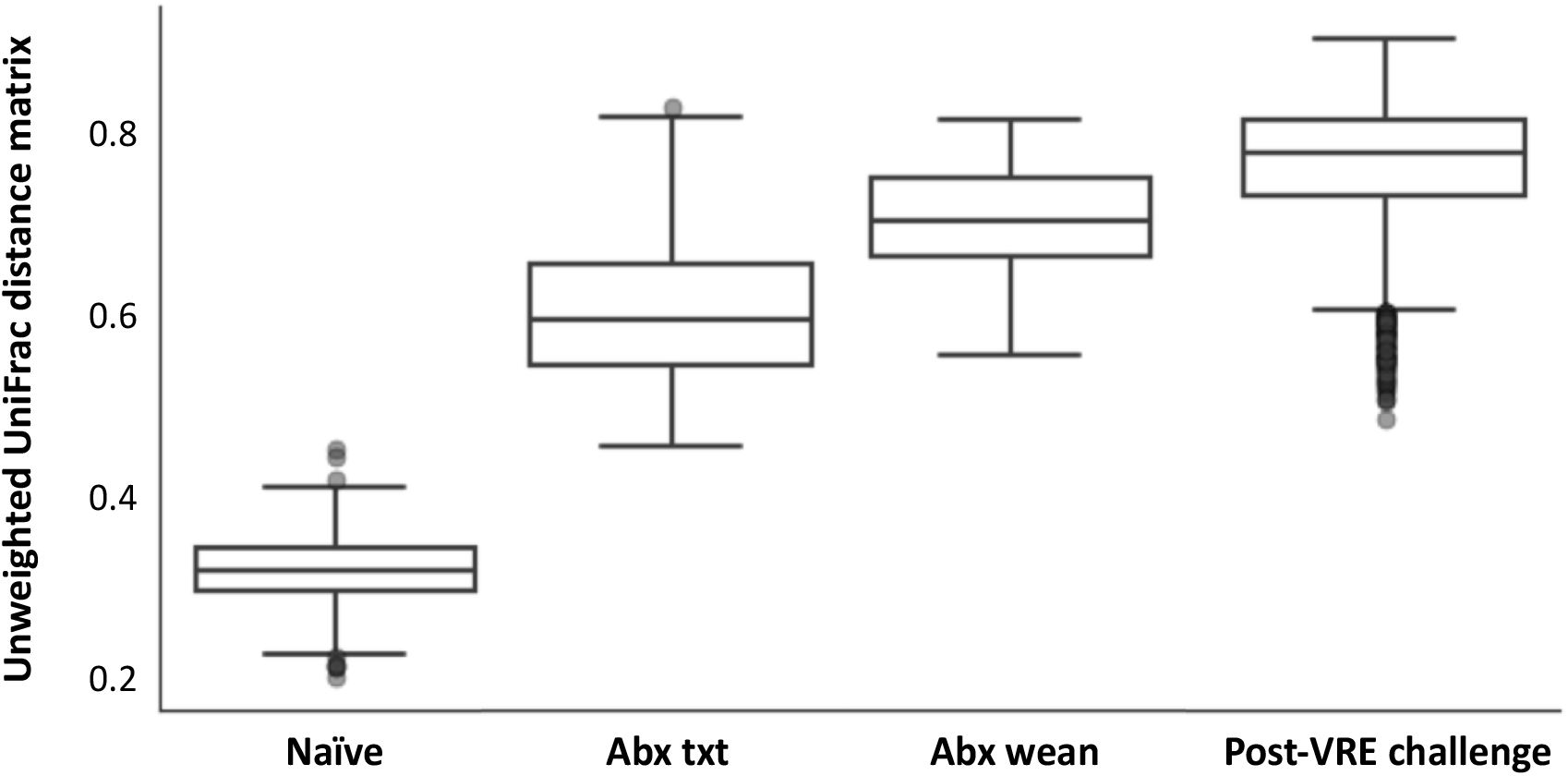
PERMANOVA testing (999 permutations) on unweighted UniFrac distances (16S rRNA gene data) relative to samples from the naïve phase. P-values for pairwise PERMANOVA testing given in parentheses for the following phases of the experiment: naïve – antibiotic treatment (0.001); naïve – antibiotic treatment (0.001); naïve – infection (0.001); antibiotic treatment – antibiotic wean (0.025).

**Supplementary Figure 3.**
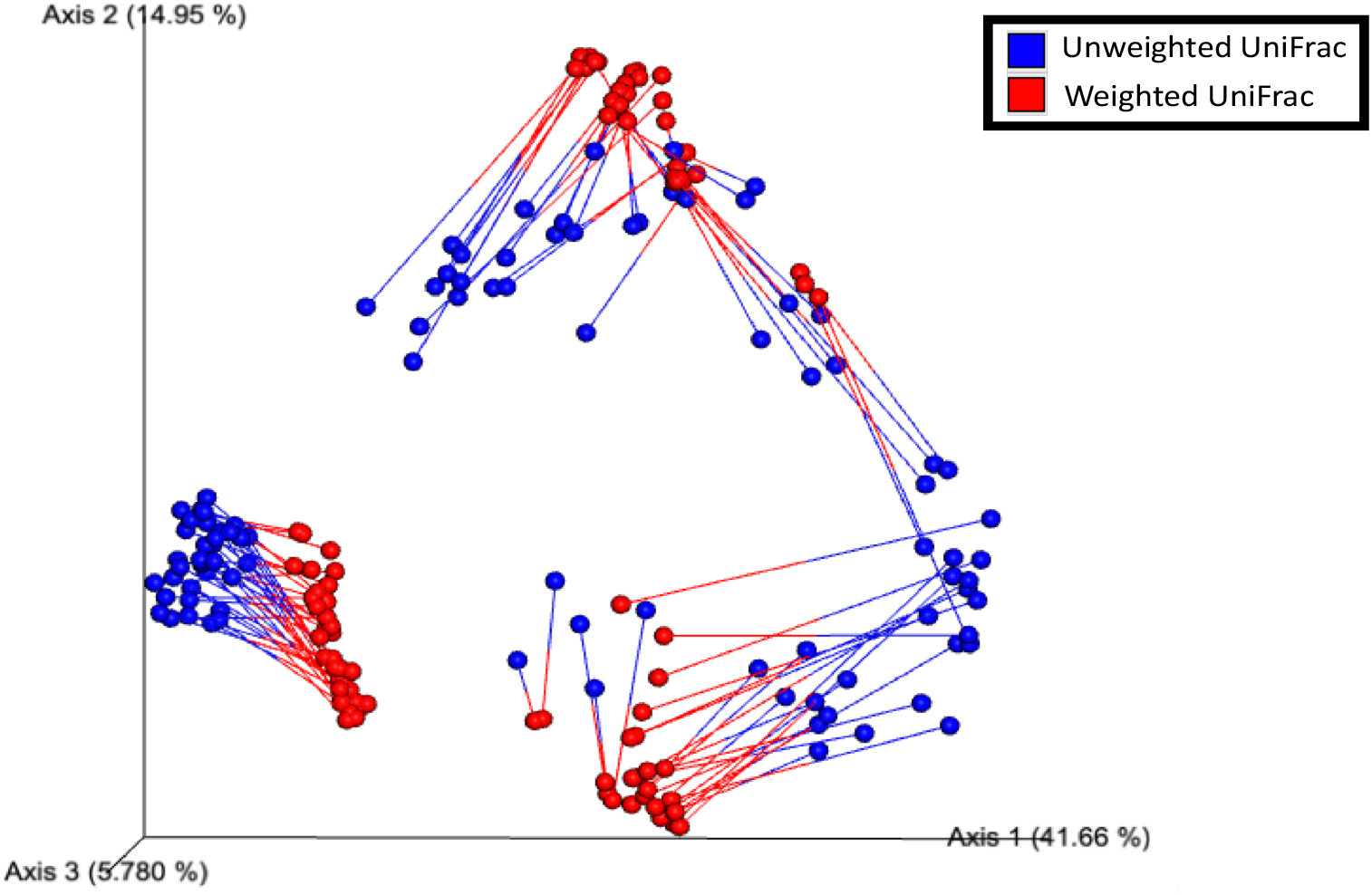
Emperor plot of procrustes analysis of unweighted (blue) and weighted (red) UniFrac distance metrices.

**Supplementary Figure 4.**
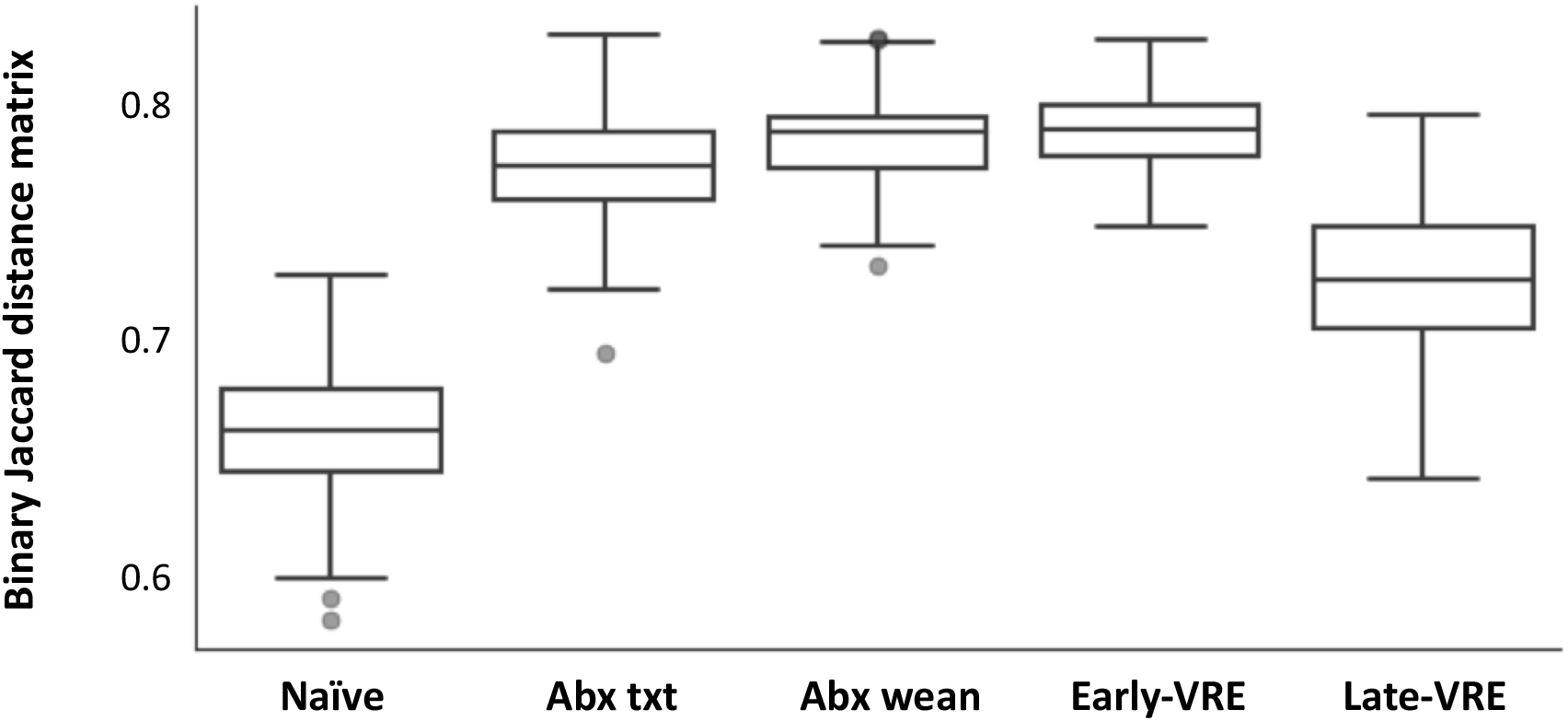
Pairwise PERMANOVA testing (999 permutations) on binary Jaccard distances (metabolome data) relative to samples from the naïve phase. While naïve and late-VRE samples are significantly different, late-VRE has a lower distance to naïve compared to Abx txt, Abx wean and early-VRE samples.

**Supplementary Figure 5.**
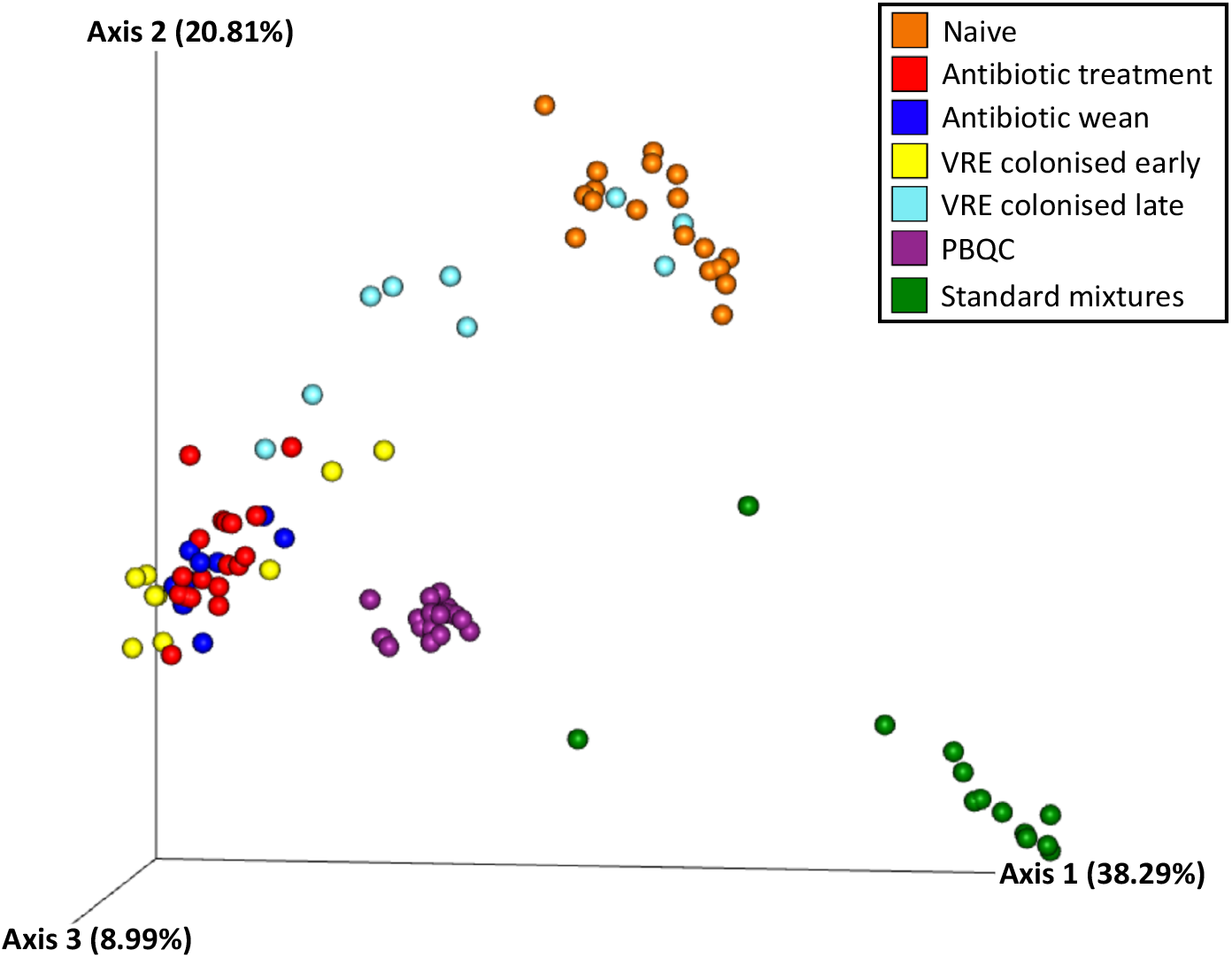
Emperor visualization displaying samples assayed for metabolomics. Principal coordinate analysis plot of distances between each sample based on their overlapping molecules as measured by Binary Jaccard. Samples assayed include pooled biological quality control samples (purple), and samples of known standard metabolite mixtures (green). The general metabolome profile of test samples is retained.

**Supplementary Figure 6.**
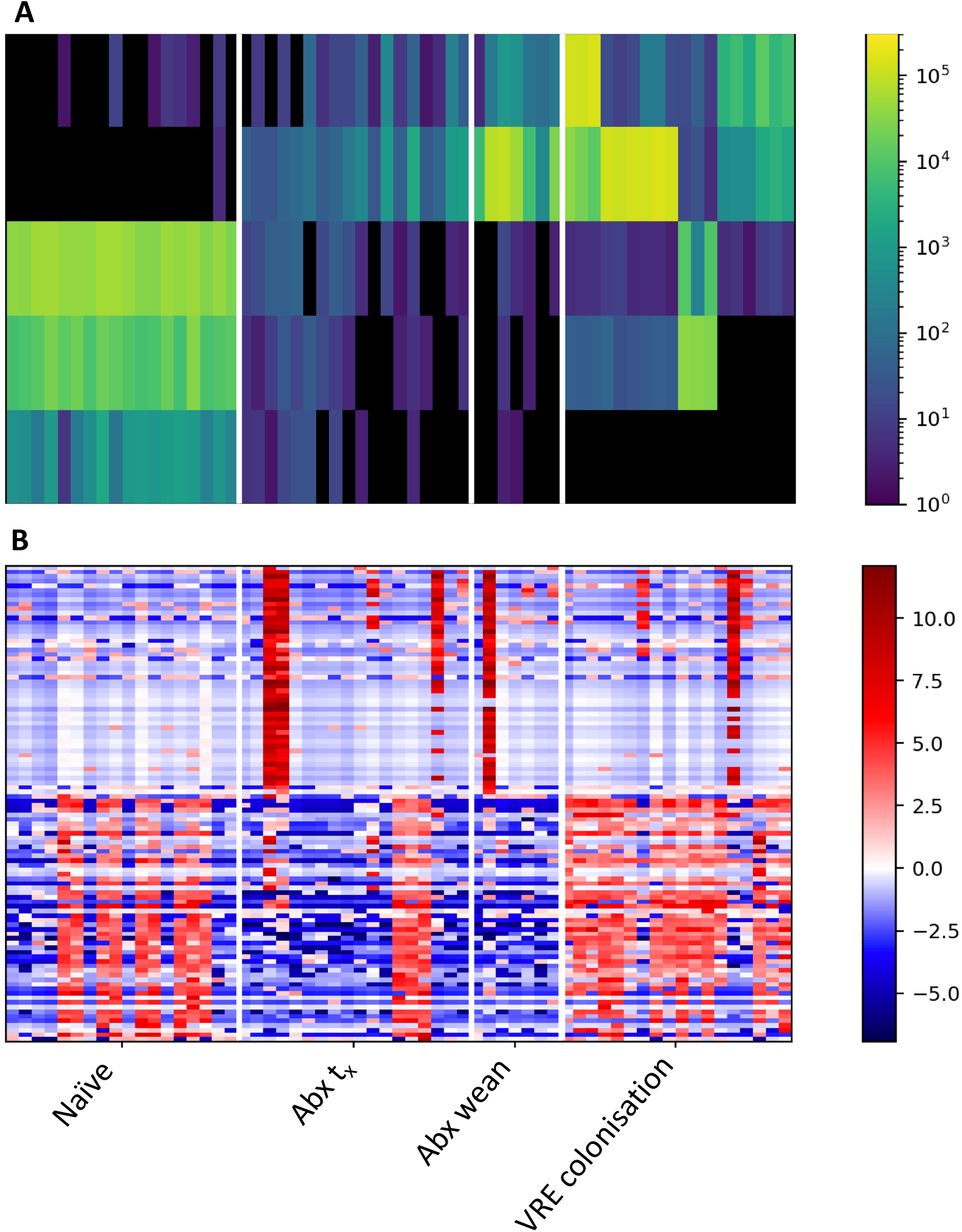
Paired feature abundance heatmaps. **(A)** Microbe abundances and **(B)** metabolite log centered abundances across each phase of the experiment. Heatmaps are aligned along the x-axis (phase of experiment); y-axis displays microbe and metabolite abundances.

